# Insights into Health Data Science Education: A Qualitative Content Analysis

**DOI:** 10.1101/2024.09.23.614482

**Authors:** Narjes Rohani, Michael Gallagher, Kobi Gal, Areti Manataki

## Abstract

**Background:** Early career researchers in Health Data Science (HDS) struggle to effectively manage their learning process due to the novel and interdisciplinary nature of this field. To date, there is limited understanding about learning strategies in health data science. Therefore, we aim to uncover learning strategies that early career researchers employ to address their educational challenges, as well as shed light on their preferences regarding HDS teaching approach and course design.

**Method:** In this study, we conducted a qualitative content analysis through semistructured interviews with ten early career researchers, including individuals pursuing master’s, PhD, and postdoctoral research programmes in HDS, across two higher education institutions in the United Kingdom. Interviews were carried out in person from June 2023 to August 2023. Data were analysed qualitatively using NVivo software. Descriptive statistics were employed for quantitative analysis.

**Results:** Regarding learning strategies, we identified ten main categories with 22 codes, including collaboration, information seeking, active learning, focus granularity, elaboration, organisation, order granularity, goal orientation, reviewing, and deep learning strategies. Regarding course design and teaching, we discovered four categories with 14 codes, including course materials, duration and complexity, online discussion, and teaching approaches.

**Conclusions:** Early career researchers used a range of learning strategies aligned with well-established learning theories, such as peer learning, information seeking, and active learning. It is also evident that learners in HDS favour interactive courses that provide them with hands-on experience and interactive discussion. The insights derived from our findings can enhance the quality of education in HDS.

## 1 Introduction

There is an urgent need to educate professional researchers in Health Data Science (HDS) [1]. Research has shown that early career researchers in HDS struggle to manage their learning and overcome their educational challenges, such as lack of foundational knowledge and difficulty of communication in an interdisciplinary environment [2]. Challenges in HDS education are not limited to early career researchers but also affect educators and institutes around the world when teaching HDS topics to learners from diverse academic backgrounds [2].

Health data science is defined as the use of computational techniques to analyse health/biological/medical data in order to discover new insights, connect those data, and describe the findings to biologists, informaticians, and other stakeholders [3]. In other words, we consider HDS as a field utilising computational methods to analyse biological, health, and/or medical data. Applying this definition, disciplines such as bioinformatics, biostatistics, and medical informatics can be considered as part of HDS, as they analyse data to address questions in biology, health, and/or medicine [2, 3].

Several recent studies [1, 2, 4–7] have emphasised the importance of providing insights into HDS education and potential strategies that can address aforementioned educational challenges. In a recent study [2], researchers identified seven grand challenges in bioinformatics, ranging from those that affect educators to those affecting learners. They emphasised conducting research to find solutions for those challenges and improving education in bioinformatics. A survey conducted by Magana et al. [8] aimed to summarise the characteristics of existing educational programmes in bioinformatics. Their objective was to inform the current state of education and training in this field. They encouraged educators to share their experiences and challenges through academic papers. They also invited educational researchers to collaborate with educators to improve bioinformatics education. The survey revealed that bioinformatics programmes mostly focused on training through problem-solving methods and were based on collaborative learning. Additionally, diverse multimedia and delivery methods were used, such as face to face, web-based learning and distance learning.

Identifying learning strategies in HDS can be useful to assist both learners and educators [9]. Previous work in the field of education ha s demonstrated the significant impact of learning strategies (approaches that someone employs to manage their learning process) on learners’ success [9–11]. Learning strategies can vary depending on the nature of the field; therefore, learners in each field may use different learning strategies [9, 12, 13]. For example, Bickerdike *et al.* [13] analysed self-declared learning strategies of undergraduate medical students in year 2 and in the final year of their study programme. They found that medical students used deep approach, surface approach, organisation, and monitoring strategies. Among the strategies identified, organisation strategy had the highest correlation with students’ performance. In another study, Matcha *et al.* [9] used data mining to identify students’ learning strategies in biology, computer engineering, and introduction to programming courses. They showed that the learning strategies used in each discipline and course are different.

Furthermore, research has demonstrated that the teaching approaches and course designs that are effective for each discipline can differ significantly [14–16]. According to Cameron [17] teachers in all disciplines are still influenced by bias and that is why they prefer to employ traditional teaching approaches, which can negatively influence student engagement. Stark [18] highlighted that teacher beliefs about educational purpose, as well as their opinions regarding the nature of their field can influence course design. Educational research [16, 19, 20] has also emphasised the importance of considering student characteristics, course topics, learning environment, and student level in course design and choosing teaching approach. Accordingly, the teaching approach and course design should be tailored to the characteristics of each discipline and its students.

There is a lack of research that sheds light on learning strategies that early career researchers in HDS use to overcome their educational challenges [2]. Also, educators and teachers do not have enough information about effective teaching approaches and course designs for HDS. The aim of this study is to address these gaps by exploring early career researchers’ learning strategies as well as by identifying effective teaching approaches and course design elements suggested by early career researchers. To the best of our knowledge, it is the first qualitative analysis that investigates learning strategies and areas to improve courses in HDS.

We conducted a qualitative content analysis by interviewing early career researchers, including postgraduate students and postdoctoral researchers in HDS, to answer the following research questions:

1. What learning strategies do/did early career researchers employ to overcome their educational challenges in HDS?
2. What teaching approach and course design do early career researchers prefer for their HDS learning?

The results show that the top three learning strategies employed by early career researchers were peer learning and help seeking, information seeking, and active learning. Using peer learning and help seeking, early career researchers communicated with their peers through online discussion forums or sought guidance from their supervisors. Information seeking involves participation in online courses, conferences, and reading books to get familiar with the field and address the lack of foundational knowledge. Active learning defined as actively engaging in problem-solving by practicing coding and studying examples. Regarding teaching approach and course design, early career researchers highlighted the benefits of teaching through discussion and problem -solving rather than passive lectures. This study provides valuable insights into learning strategies and effective teaching approaches and course design in HDS that can help address challenges in HDS education.

## 2 Methods

In this study, the instructions of the Standards for Reporting Qualitative Research (SRQR) [21] were used to report the methodology of our qualitative analysis. These guidelines comprise 21 criteria aimed at enhancing transparency in reporting qualitative research (the SRQR checklist is available in **Additional file 1**). By adhering to these guidelines, we ensured that participant information, researcher characteristics, and detailed explanations of data analysis procedures were described comprehensively.

### 2.1 Study Design

The present study was a qualitative content analysis, involving semi-structured interviews with early career researchers, including postgraduate students and postdoctoral researchers in the field of health data science in two higher education institutions in United Kingdom (UK).

Our objective was to answer the specific research questions in order to gain insights into the challenges, learning strategies, and course preferences of early career researchers, studying and working in the HDS field. We utilised prior knowledge derived from literature and our quantitative analysis to formulate our research questions, frame the interview questions, and categorise quotes using qualitative content analysis. Although our prior research findings [22] were used to guide analysis, there was also the potential for revealing additional knowledge.

In other words, this study was carried out from post-positivist point of view [23] using conventional inductive qualitative content analysis [24]. We chose post-positivist research paradigm because this approach acknowledges the complexities and biases of the social aspects while still valuing empirical evidence. The traditional approach of inductive content analysis is typically suitable when there are limited prior studies or grounded theories outlining the phenomenon under exploration, or when the phenomenon itself is not well-defined [24, 25]. Consequently, this study does not adhere to a single strict theory. Nonetheless, considering the study’s aims and the observed categories and subcategories, some of our findings can be interpreted through the lenses of cognitive and metacognitive theories of self-regulated learning [26, 27], Bigg’s learning theory [28], and the MSLQ theory [11].

We started the interview with asking participants about their research study and then asked them a series of more specific questions (the list of questions is available in **Additional file 2**) related to the research questions. We continued the data collection until coding saturation was achieved; the prevalence of new codes significantly diminished. The format of the interviews was one to one and in person in a private meeting room. The interviews were conducted from June 2023 to August 2023, with each interview lasting between 15 to 30 minutes, which proved sufficient given the specific focus of the study.

Ethical approval for this study was granted by the School of Informatics, University of Edinburgh. All interviews were recorded in audio format and subsequently transformed into anonymised interview transcriptions using customised Python code, with each participant assigned a unique code and their personal information removed from the data. The audio files were deleted after transcription. Prior to the interviews, participants received a participant information form, containing the research goals, the interview procedure, and ethical considerations. During the interviews, participants declared their consent through signing the consent forms (sample participant information and consent forms are available in **Additional file 3**). To ensure secure storage, all consent forms and associated files were stored in a password-protected computer, which is only accessible to the researchers.

### 2.2 Participants

We initiated the recruitment process by sending invitation emails to early career researchers in HDS across two higher education institutes in the United Kingdom. The sampling strategy was purposeful, taking into account the fact that those invited to participate are early-career researchers in HDS. Our recruitment process was carefully designed in a way to ensure having a suitably diverse data set that is essential for qualitative analysis. The following factors were taken into consideration:

- Inclusion of participants with diverse backgrounds, including both participants with biological/health/medical and computational academic backgrounds.
- Incorporation of participants from a wide range of ethnicities.
- Selection of two institutions situated in different cities within the UK to capture regional variations.
- Involvement of participants in different academic levels, including master’s, PhD, and postdoctoral.
- Ensuring a balanced gender distribution among the participants.

Once the prevalence of new codes greatly diminished, we concluded the data collection. We successfully recruited a total of 10 participants, which was deemed to be a sufficient sample size given the specific focus of this study. There were no exclusions, and all early-career researchers who volunteered were included in the study. Among these 10 participants, 6 were affiliated with one institute, while the remaining 4 participants were from the other institute. Gender distribution was evenly split, with half of the participants being female and the other half male. The participants’ age ranged from 20 to 35, originating from diverse regions such as Asia, Europe, the Middle East, and South America.

Regarding education level, 6 participants were PhD students, 2 were postdoctoral researchers, and the remaining 2 were master’s students (one of them recently finished their master’s program and one were at the final year of their program). In terms of academic backgrounds, 6 participants had an academic background in computational fields such as computer science, while the remaining 4 had biological backgrounds such as cell biology. Notably, among those with biological backgrounds, half declared a basic knowledge of computational concepts (mixed).

### 2.3 Researcher characteristics and reflexivity

All the interviews were carried out by lead author, NR, who is a PhD candidate in Precision Medicine from a different institution (University of Edinburgh) than the participants. NR has an academic background in computer science during her master’s and undergraduate degrees. Before conducting the interviews, NR learned about how to conduct qualitative data analysis and made sure to follow unbiased data collection and interpretation. NR had no personal or professional relationships with the participants prior to the interviews, which helped ensure unbiased data collection process. To make sure the interviews were unbiased, NR practiced self -reflection and took notes to be aware of any potential biases. NR also considered ethical aspects during the interviews to minimise any possible biases. The generated codes and findings were reviewed by and edited with the help of all the authors in order to discuss any potential biases in the interpretation.

### 2.4 Data processing

A custom Python script that uses AssemblyAI’s Python SDK [29] was employed to transform the audio-recorded interviews into text forma t. Afterwards, the lead author read and checked the transcribed interviews to ensure accuracy of transformation, and she applied any changes required. Finally, the transcribed interviews were imported into NVivo 14.23.2 software [30], which was utilised for coding the transcribed interviews. We started the coding process after the initial interview, and we carried out this process in a recursive way. During coding, NR and MG collaborated to refine the codes until achieving agreement and data saturation.

## 3 Synthesis and Interpretation

In this section, we present the results of our analysis across our two research questions regarding learning strategies in HDS as well as teaching and course preferences. Each category and code were supported with quoted evidence and supplemented by relevant well-known learning theories.

### 3.1 What learning strategies do/did early career researchers employ to overcome their educational challenges in HDS?

We asked participants how they managed their educational challenges in HDS. Ten categories with 22 codes emerged. In the rest of this section, we will discuss each learning strategy (categories and codes) as mentioned by the participants. Error! Reference source not found. shows a summary of the learning strategies identified.

#### S1: Collaboration

The collaboration category (30 references by all 10 participants) was frequently discussed by participants. This category included five codes: **peer learning and helpseeking** (24 references by 10 participants), f**lexibility in solo and group learning** (5 references by 5 participants), i**nterdisciplinary group** (4 references by 4 participants), and **independent learning** (1 reference by one participant).

Regarding **independent learning**, only one participant who was a PhD student with mixed background declared that they prefer to work independently when working on a project:

> *“[..] If you want to learn the most in the shortest available time, I think it has to be independent (P9).”*

However, half of the students (5 of 10) discussed f**lexibility in solo and group learning**, indicating that it depended on factors, such as the project’s nature, group members, and other considerations as these factors also mentioned in previous study [31]. For example, a PhD student with computational background mentioned:

> *“It depends on the project because it means you are not a master of everything. For example, if I am pursuing a project in a field, I’m knowledgeable in, I may choose to work independently. However, if it’s in an area where I lack expertise, I might opt fo r teamwork (P4).”*

Four participants also discussed the advantages of having an **interdisciplinary group** that includes members with diverse backgrounds. They explained that this diversity can facilitate constructive discussions and collaboration. For instance, one PhD student with computational background highlighted,

> *“I mean for multi-discipline project it’s better to have work on the group because for example my expertise is about machine learning and other expertise may be about interpretation so that’s why even now in this group we work together as a team (P3).”*

The final and highly referenced code is **help-seeking and peer learning** (referenced 24 times by all participants), which includes references related to learning through group discussions, collaborative team projects, and seeking guidance from others. This category is aligned with the well-established learning theories, such as self-regulation learning, and peer learning [11, 27, 32]. For example, one postdoctoral researcher with computational background reflected on their learning strategy as group study during their PhD:

> *“Me and my colleagues, my classmates, we had our study groups that we gathered together and studied those books together in order to get to a point to understand those biological courses. That was the first thing. The second thing, […] we also gathered together and discussed the materials together. And those kinds of discussion really helped me. (P1).”*

Participants also discussed that collaboration and help-seeking strategies should not necessarily be limited to peers and supervisors. Online discussion forums, such as Stack Overflow can be used to ask questions. For example, a PhD student with computational background mentioned:

> *“There are number of discussion forums available for bioinformatics and also stack overflow and something like that. So that is really good because if you are stuck with any implementation something like that, that is going to help you a lot. So I think di scussion forum is a really good for researchers here (P4).”*

In summary, the collaboration learning strategy was discussed 30 times by all learners and it is the most frequently discussed learning strategy. This category includes four codes: peer learning and help-seeking, interdisciplinary team, flexibility in working as a team and alone, and working alone. The results suggest that early-career researchers in HDS find help-seeking strategies and peer learning strategies useful, and the majority of them are positive about working in a team. However, various factors, including the academic level of group members, project type, time constraints, and other considerations, may have influenced their choices. [31]. Nevertheless, it is evident that learners seek assistance by interacting with colleagues, peers, or utilising discussion forums.

#### S2: Information seeking

The information-seeking category (referenced 21 times by 9 participants) encompasses three codes: **building foundational knowledge** (referenced 9 times by 7 participants), **reading articles and participating in online courses** (referenced 8 times by 6 participants), and **familiarisation** (referenced 4 times by 3 participants). This strategy involves dynamic steps taken by learners to acquire additional information from various resources, such as papers, books, courses, and more. This strategy is consistent with self-regulation learning theory [32, 33].

**Building foundational knowledge** (referenced 9 times by 7 participants) related to quotes where learners express their approach to acquire essential basic knowledge. For example, a postdoctoral researcher with a computational background explained their strategy for gaining fundamental biological knowledge during their PhD:

> *“During my PhD, the courses that I took were really hard. At the very first sessions, I remember I had a lot of headaches. What I did was I borrowed some high school books of biology from people who were studying biology in the high school. And I first read those to get a very basic understanding of what, for example, DNA is, what RNA is, and what that sort of things, to then understand what, for example, metabolism is. And that really helped me (P1).”*

The next category relates to references concerning participation in **online courses and conferences or reading articles and books courses** (referenced 8 times by 6 participants) to enhance their knowledge but not limited to foundational knowledge. For instance, a PhD student highlighted:

> *“I improved my learning by joining the different courses with the different sites like the Udemy is there, Coursera is there then at the same time reading different general papers, conference paper, books (P4).”*

The final code, **familiarisation** (referenced 4 times by 3 participants), describes strategy in which learners familiarise themselves with a course, field, or concepts. For example, a PhD student emphasised:

> *“Mostly in the cases that I have absolutely no idea about the content of the course. I play a little bit of all of the parts of the courses to just make myself familiar with the terms that use in this course, to make myself familiar with the atmosphere, with the approach that the person who is teaching is using (P2).”*

In summary, seeking information was one of the frequent strategies discussed by the participants (referenced 21 times by 9 participants). It can be concluded that early career researchers in HDS endeavoured to acquire foundational knowledge in their field, which may be lacking due to the interdisciplinary nature of the health data science [2]. Additionally, they employed the familiarisation strategy to acquaint themselves with terminology, courses, and the field before studying them deeper. To achieve these goals, they utilised diverse resources such as online courses, articles, books, and more.

#### S3: Active learning

The **Active Learning** category contains three distinct codes: r**ehearsal in coding**, **sample-based learning**, and p**roblem solving**. Active learning is defined as an approach that emphasises learning through engagement, such as solving an actual problem to learn [34]. Among all 10 participants, 6 of them (60% of participants) referred to this learning strategy 17 times.

The first code is **rehearsal in programming** (9 references by 4 participants), which as emphasised by participants as a fundamental strategy for their learning process. A master’s student with biology background mentioned employing rehearsal in programming as an effective learning strategy:

> *“[…] I think my strategy is really just to practice coding (P10).”*

This strategy was not only effective for master’s students but also a PhD student with biology background similarly mentioned:

> *“[…] I mean, all I really did was practice, […] but to be honest, lecture wasn’t as useful as just practicing (P9).”*

Although rehearsal in programming was employed by all learners in different level of study, including postdoctorate, PhD, and master’s degree, this strategy was used more by participants who have biological (4 of 9 references) or mixed backgrounds (4 of 9 references) rather than computational (1 of 9 references). This is expected, as learners who have computational background have stronger knowledge in programming and computational aspects and these aspects are less challenging for them.

In the next code, **sample-based learning** (5 references by 3 participants), participants discussed the value of looking at examples and sample projects to learn. For example, one master’s student with biology background highlighted the benefit of looking at scripts written by others in other to learn about programming:

> *“I think it’s very helpful to look at other people’s code and try and understand what they’re doing (P10).”*

Similar to the rehearsal in programming strategy, this strategy also was discussed more by participants with biological background (4 of 5 references) compared to computational one (1 of 5 references).

In the final code which is the **problem-solving** (3 references by 3 participants), participants talked about hands-on experiences, actively engaging with real-world problems instead of passively acquiring knowledge [35, 36]. For example, one PhD student with computational background reflected on their strategy in starting learning by having a project and learning through the journey of solving that:

> *“For me, I tend to do learning by doing, so what I mean is I start with a project and then if I don’t know anything, I will start to look up […] to that particular one (P3).”*

Converse to rehearsal in programming and sample-based learning, problem-solving learning strategy was discussed only by participants with computational (2 of 3 references) or mixed (1 of 3 references) academic backgrounds.

In summary, the active learning strategy was referenced by 6 participants 17 times. This learning strategy encompasses three codes: rehearsal in programming, sample-based learning, and problem-solving. According to results, participants with a biological background employed rehearsal in programming to improve their computationsl skills. They also referred more often than participants with a computational background to sample-based learning. Through sample-based learning, they attempted to view sample codes to actively learn how to write or modify code. Conversely, problem -solving, where participants prefer to have a project and actual problem and learn through the journey of solving it, was discussed more by participants with a computational background.

#### S4: Organisation

According to the self-regulation learning theory and Motivated Strategies for Learning Questionnaire (MSLQ) learning theory [11], the organisation learning strategy involves approaches that learners use to structure and manage their learning process [11, 32, 37]. This strategy is focused on improving how learners arrange information, resources, and their study routines [11, 32, 37]. This category (referenced 16 times by 7 participants) has three codes: **time management** (referenced 7 times by 4 participants), l**ooking at the course syllabus** (referenced 5 times by 4 participants), and **prioritising** (referenced 3 times by 3 participants).

The first code is **time management** (referenced 7 times by 4 participants), which reflects the learners’ strategies in managing their time and having organised time allocated to learning and research. For example, a PhD student said:

> *“Nowadays the life is very fast. So, people do not have that much of time. So, my preference is that if I get the complete knowledge in particular videos or book or something like that, that will help me to save my time (P4).”*

The strategy of **looking at the course syllabus** (referenced 5 times by 4 participants) can help learners decide which topics are more interesting to them, what topics are more important to focus on, or familiarise themselves with a course. For example, a PhD student discussed checking the course syllabus to determine if the course topics are interesting:

> *“Actually, the first thing that I do when I face a new course, I check the topics that they say that they will teach in this course. I will check the topics to see that whether those topics are interesting or not (P2).”*

Participants also used the **prioritisation** (referenced 3 times by 3 participants) strategy when reading an article, taking a course, and/or working on a project. For example, a postdoctoral researcher said:

> *“I prioritised the concepts based on what is needed for the project and the ones that are not relevant much (P1).”*

Interestingly, organisation strategies were discussed more by PhD students (11 of 16 references). It can be due to fact that they have hectic schedules, and it makes sense that they are concerned about time management and prioritise important topics to manage their time and effort. Also, according to results, these strategies were more employed by early career researchers with computational background (11 of 16 references).

In summary, the organisation strategy is referenced 16 times by 7 participants. This indicates that organisation is an important learning strategy for postgraduate students, as it helps them prioritise their learning by focusing on necessary concepts for their research and managing their time in a way that allows them to progress with their projects on time.

#### S5: Order granularity: sequential vs global

The **sequential** learning strategy (referenced 12 times by 7 participants) refers to an approach in which learners follow learning topics in a step-by-step order, based on a predetermined structure [38–40]. This strategy demonstrates that learners adopt an organised and linear approach to navigating topics. On the other hand, in the **global learning** strategy (referenced 3 times by 3 participants), learners engage with learning materials in a non-sequential and unpredictable way. Unlike sequential learning, where learners adhere to a predetermined structure, global learning might jump between different topics or sections [38–40].

For example, a master’s student discussed the advantages of a **sequential** educational program:

> *“It did help that my master’s course was very sequential, where they started off with the basics of coding and along that, they also taught the basics of statistics, and they also provided extra classes beyond the core curriculum so that you can kind of learn the coding more in-depth and be more prepared for the courses as well (P7).”*

In contrast, a few participants mentioned employing the **global learning** strategy. For instance, a PhD student noted:

> *“I would say jumping between different concepts works for me […] (P9).”*

In summary, the majority of participants (7 out of 10 in total) mentioned sequential learning as their strategy over global one. They discussed how having a step-by-step guideline can be useful, suggesting that a sequential course design and clear step -by-step instructions could enhance their learning experience.

#### S6: Elaboration strategy

The elaboration category (referenced 11 times by 7 participants) includes three codes: **notetaking** (referenced 7 times by 6 participants), k**nowledge integration** (referenced 3 times by 3 participants), and **summarisation** (referenced once by one participant). The elaboration strategy, consistent with the MSLQ [11], assists students in encoding information for long-term retention by establishing internal connections among concepts [11]. This strategy encompasses paraphrasing, summarisation, and generative notetaking, enabling learners to merge and relate new information with their prior knowledge [11].

**Notetaking**, referenced 7 times by 6 participants, was discussed as the most employed elaboration learning strategy. For example, one participant mentioned notetaking as an effective approach to converting information from research papers into understandable information that they can easily refer to later.

> *“I take note, yes. Now, you always have your notes with you. You always try to transform a research paper to a more, we can say, figures and that is easier to keep a track on what you learned, because it’s not easy to learn something from scratch. […] (P5).”*

The **knowledge integration** strategy (referenced 3 times by 3 participants), where participants integrate new information with their prior knowledge for better understanding, was discussed 3 times by 3 participants. For example, a researcher that recently finished their master’s degree reflected on using the knowledge integration strategy during their master’s program:

> *“My course included machine learning modules, and prior to enrolling in my master’s course, I had some basic experience with machine learning. To enhance my understanding, I refer to what I’ve previously written in my work. While it may not always be directly relevant to my current coursework, I’ve found that reviewing my previous notes is helpful (P7).”*

The same researcher also discussed the **summarisation strategy**:

> *“And also, I try to summarise it in a more concise manner so that I can find the content that I need more easily as compared to looking through the entire presentation they have on the content (P7).”*

In summary, the elaboration learning strategy, which involves facilitating the learning of new concepts and their retention in long-term memory by connecting them to prior knowledge, taking notes, or summarising, was discussed by 7 participants a total of 11 times. Participants from all degree levels and academic backgrounds have employed this strategy. Among these elaboration strategies, notetaking was employed most frequently (7 times by 6 participants).

#### S7: Focus granularity: details vs general information

This learning strategy (referenced 5 times by 4 participants) shows whether participants choose to concentrate on **detailed** information or opt for a focus on more **general** concepts and ideas within a given subject or topic [38]. One learner mentioned focus on details in the subject and paying meticulous attention to specific facts and theories (referenced once). Conversely, other learners prioritise general concepts (referenced 4 times by 3 participants) by emphasising main and general ideas. They prefer to concentrate on high-level ideas and may skip or briefly skim through detailed content [38].

For instance, a PhD student discussed their preference for focusing on the main objective initially and studying the details related to the project later:

> *“When started working on such projects, especially in areas like AI, machine learning, or data science related to cancer research, it’s better to begin with a data-driven approach rather than a biology-driven one because that will drive you crazy. Once you have gone overwhelmed, you cannot move further. So, when you start from a data - driven approach and then you start making some sense, you can focus on details in your research interest. And then you try to learn those detailed terminologies (P6).”*

Conversely, another PhD student declared a preference for courses rich in details, highlighting their interest in using detail focus strategy:

> *“So definitely you need to provide such a course which is giving the detailed information about each and everything with the practical and with theory and explaining each and everything (P4).”*

In summary, focus on general information (referenced 4 times by 3 participants) was more popular than focus on detail information (referenced once by one participant). It can be inferred that main ideas were important for early career researchers in health data science. They preferred not to allocate time to less important details during their initial learning stages. They decided to acquire detailed information while actively working on their projects. Therefore, it might be beneficial to place greater emphasis on main ideas in both teaching and course content, with detailed information servin g as supplementary material within the course.

#### S8: Goal orientation

Goal orientation category (referenced 4 times by 3 participants) is consistent with the strategies defined in self-regulation learning theory and MSLQ [11, 32]. By using the goal orientation learning strategy, learners establish specific learning goals, which can encompass both short-term and long-term objectives [11, 32]. Their subsequent efforts and actions are guided by these goals, whichshows the importance of a clear sense of purpose in the learning process [11, 32]. Learners who adopt goal orientation tend to exhibit greater curiosity and motivation [11, 32]. This category comprises two codes: **making a plan** (referenced 2 times by 2 participants) and t**aking extra classes beyond the core curriculum** (referenced 2 times by 2 participants).

Making a plan refers to creating structured and organised schedules for studying or preparing for assessments. For instance, a researcher with master’s degree and mixed background emphasised their plan-making approach for assessment, stating:

> *“If it’s coding-related assessments, I usually start with doing a plan of what is the process (P7).”*

There were early career researchers that aim to extend their learning beyond the course syllabus by seeking out extra classes. For example, a PhD student highlighted the importance of extra study, saying:

> *“[…] we should be self-learners. So, we should not be restricted with what we learned from the lectures. We should go beyond that (P6).”*

In summary, goal orientation learning strategy was discussed by 3 early career researchers 4 times. The results show that early career researchers employed the planning and going beyond curriculum as goal orientation learning strategies. This result is in line with previous studies that show that learners with clearly defined learning goals tend to be more focused and driven, often displaying a greater desire for in -depth knowledge in their field [11, 26, 32]. Consequently, they are more inclined to engage in activities such as taking extra classes and making plan for their learning journey [11, 32].

#### S9: Reviewing learning materials

The reviewing learning strategy (referenced 4 times by 3 participants) involves revisiting and studying previously learned materials, such as lecture notes, books, recorded video lectures, or other educational resources. Based on self -regulation learning theory, this strategy is used by learners to reinforce their knowledge and refresh their memory [32, 33].

For example, a master’s student declared that they review learning materials for assessment:

> *“For assessments, I usually try to review what they taught during the lectures (P7).”*

In summary, the reviewing learning strategy, which is accordance with self-regulation learning theory, is usefulto reinforce understanding and refresh memory. However, it was discussed less frequently by early career researchers (referenced 4 times by 3 participants).

#### S10: Deep learning

This strategy (referenced 3 times by 3 participants) is about achieving a clear and thorough understanding of a concept, idea, or topic. This strategy (compatible with the deep learning strategy discussed in the Biggs theory, which is a well-recognised learning theory [41–43]) involves obtaining a comprehensive grasp of fundamental knowledge and connections between different concepts [42, 44]. This strategy was only mentioned by PhD students with computational backgrounds. For example, one of them high-lighted:

> *“When I encounter something that seems a bit unclear, I typically return to it and conduct online research, such as using Google (P5).”*

Therefore, learners who employ this strategy do not allow any concept to remain unclear to them, as they are keen to gain a deep understanding of all topics.

### What type of teaching approach and course design do early career researchers prefer for their HDS learning?

We asked participants about their preferences regarding effective course design and teaching approach for HDS education. Based on their answers, four categories emerged that explained their preference regarding course materials, course length and level, discussion forum, and teaching method. In the rest of this section, we discuss each category and the corresponding codes.

**Table 2** presents a summary of the findings of RQ2.

**Table 1.**
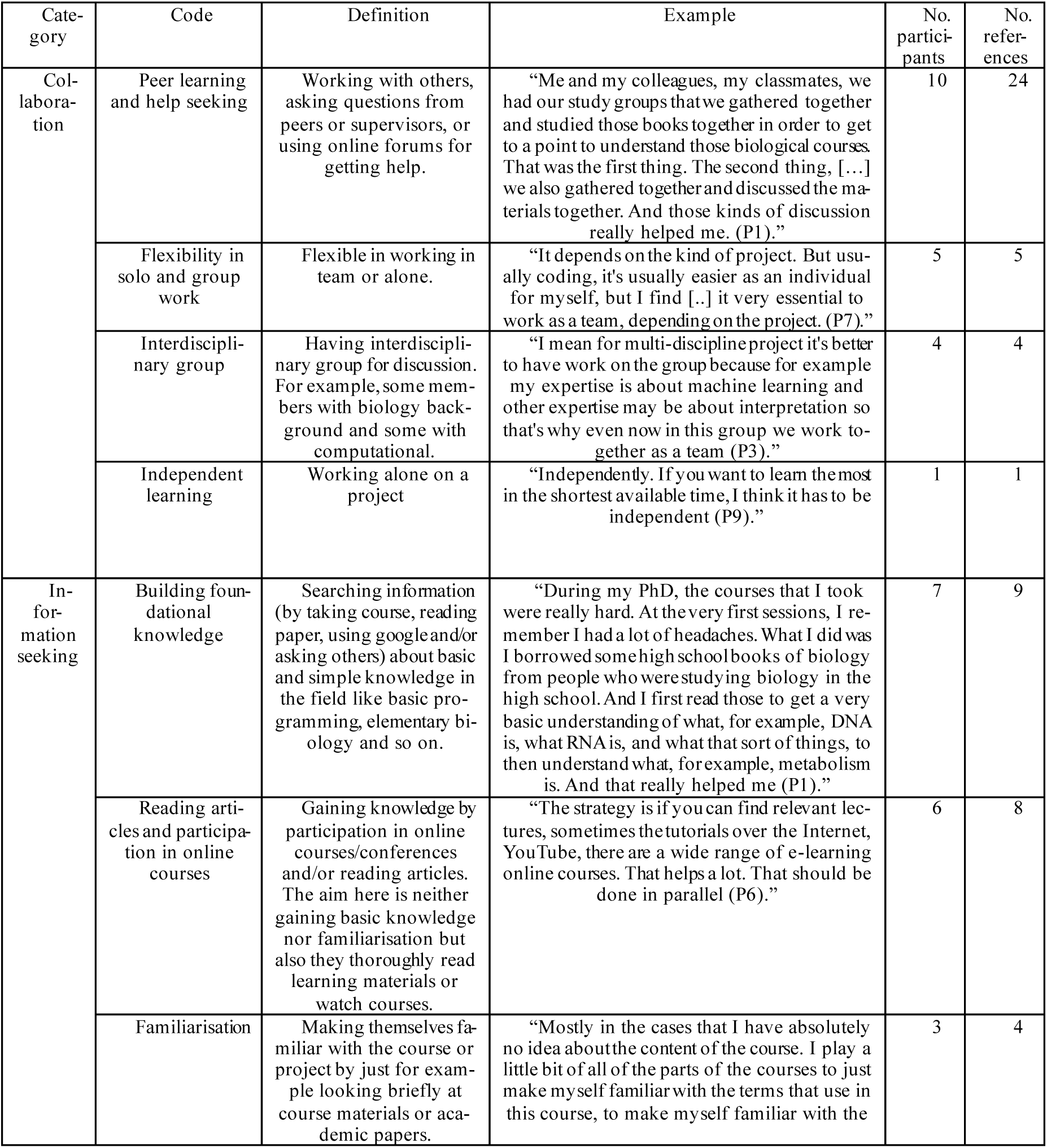
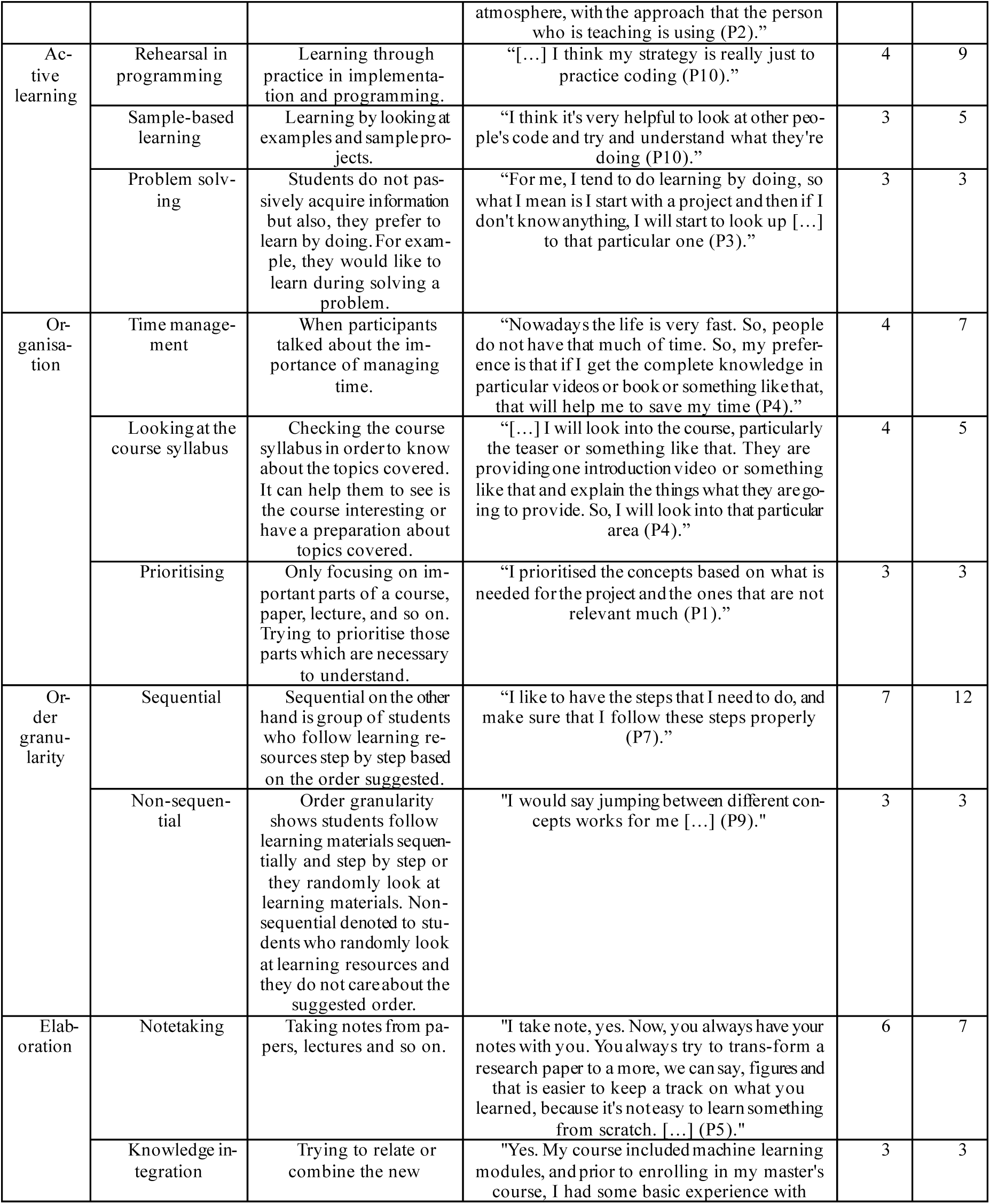
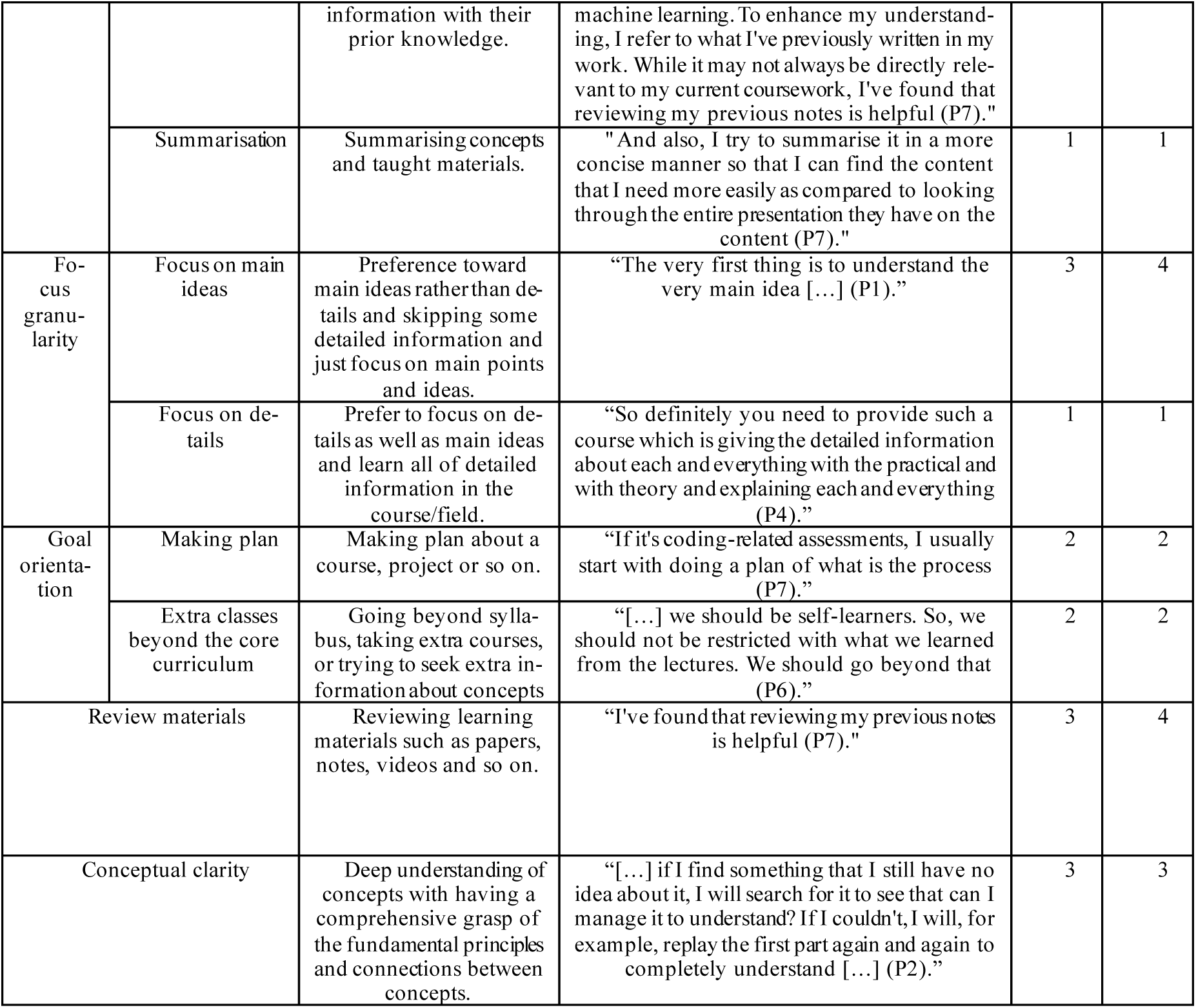
Summary table showing the results of RQ1: What learning strategies do/did early career researchers employ to overcome their educational challenges in HDS?

**Table 2.** Summary table showing the results of RQ2: What teaching approach and course design do early career researchers prefer for their HDS learning?

| Category | Code | Definition | Example | No. participants | No. references |
| --- | --- | --- | --- | --- | --- |
| Course materials | Visual material | Quotes about course learning resources and contents. For example, preference toward having textual materials or video lectures. Or talking about some topics of interest to be taught. | "I prefer slides, the things that are also visual. [...] (P1)." | 6 | 8 |
|  | Reading and textual materials |  | "It depends on individual personality. Some people love to watch. Okay. For me, I love reading research papers (P6)." | 4 | 4 |
|  | Instruction in technical tools |  | "I wish apart from just the coding aspect, they also taught us how to use a HPC (High-Performance Computing), or how to use GitHub, things like that. I think that would have been very helpful (P10)." | 2 | 2 |
| Course length and level |  | Quotes about length or level of a course. | "I think I do wish that we had more, because at the start, we only had a very short course of coding. I wish that was a bit longer (P10)." | 3 | 5 |
| Discussion forum | Immediate support | Immediate answer and support in online discussion forums | "I rarely use discussion forums because when I have a question while in the middle of a course and I want to grasp a new concept, I prefer using chat GPT. It provides instant answers, unlike Google and discussion forums, where you need to search through multiple search results to find the right answer (P2)." | 6 | 7 |
|  | Engaging all students | Engaging all students in a discussion | "I mean, usually if you have an issue, you start with discussion, but sometimes people do not engage. Or maybe because I'm not particular interested in that topic. So that's the reason I feel the discussion. So, the success of online course discussions depends on the level of engagement (P3)." | 2 | 2 |
|  | Subject-based discussion | Having subject-based discussion that students know to talk about what subject. | "[...] For example, on platforms like Stack Overflow or other similar websites, discussions usually are around solving problems. If it's a common problem, discussions tend to be more active. Everyone shares their perspectives on how to solve the problem, which creates engaging discussions. However, discussing just the course material without a specific problem to solve can be challenging for students to collaborate on (P5)." | 1 | 1 |
|  | Asking basic questions | Encouraging students to ask even basic questions | "The thing is, some people might think if they ask their question, that might seem silly or very simple. But having that said that every single question is really important, even if it might seem simple, you should ask it (P1)." | 1 | 1 |
|  | Anonymised question | Option of asking question anonymously that can improve discussion | “[...] Maybe if students are shy, they can have an anonymized feature where people don't reveal who they are and they can ask a question, so they don't feel like they're asking anything stupid or something (P7).” | 1 | 1 |
| Teaching approach | Teaching through discussion | Teaching concepts through discussion | “I like it when we're all trying to work on one problem and then talking about it and solving it together rather than lecturing teaching (P10).” | 8 | 15 |
|  | Lecture before discussion | Having lecture before discussion. This method is opposite of the previous one. | “I think you need to have some theory and then apply to a practical part because otherwise it's like just theory and you don't know how to be a bit more flexible to apply and then actually come with the problem that arises when you face the situation (P8).” | 1 | 1 |
|  | Practical emphasis | Teaching by more focus on practical concepts rather than theory. | “If it is more coding or computing way of the courses, then I would suggest having hands-on experience with coding. If it is more on the biological side, I also suggest having some hands-on experience of the biological (P1).” | 8 | 17 |
|  | Theoretical emphasis | Teaching by more focus on theory rather than practice. | “I think you need to have some theory and then apply to a practical part because otherwise it's like just theory and you don't know how to be a bit more flexible to apply and then actually come with the problem that arises when you face the situation (P8).” | 2 | 2 |
|  | Teaching with simple language | Teaching technical concepts by using simple language and simplification. | “The video lectures that have very simple cartoon things in it that makes everything simpler so that all the audiences with every level can understand that [...] I should say then simplified videos (P1).” | 1 | 3 |
|  | Having interdisciplinary lecture | Teachers with variety of academic background and having an interdisciplinary lecture design | “For me if during my PhD or the first few days of the postdoc, if there was a biologist who came to our sessions and talk about that computational problem from a biologist perspective, it's like an inviting lecturer that for example, you have a lecturer that is learning you how to do the statistical analysis on the biological data. And you have, for example, twelve sessions that a computational person is teaching you. Second, you might have an introduction of how clinical people are talking, what their language is, and this can really help if it is happening in the earlier stages in their academic life (P1).” | 1 | 2 |

#### P1: Course materials

We asked participants about their preference regarding learning resources. Among their answers, three codes emerged: **visual materials** (referenced 8 times by 6 participants), **reading and textual materials** (referenced 4 times by 4 participants), **instruction in technical tools** (referenced 2 times by 2 participants).

The results show that visual resources were discussed more compared to textual materials. For example, one postdoctoral researcher highlighted:

> *“I prefer slides, the things that are also visual. […] (P1).”*

However, there are participants who discussed the benefit of reading materials. For example, one of the participants that recently graduated with master’s degree in HDS stated:

> *“I do like to read more. I like to have something written down so that I can refer back to them. I find that video lectures, they tend to miss out on a lot of details, and I want something in detail so that I can pick out the information that I want (P7).”*

Although the number of participants and references of visual materials (6 of 10) is higher than textual materials, this difference is not big. This could be due to the fact that reading research papers is a significant part of early career researchers’ study and they need to have access to details related to their research in written format; therefore, they can find those details later when they need them.

Another code involves including topics related to **technical tools** (referenced 2 times by 2 participants) and skills such as GitHub and Linux, which are not typically taught in health data science courses. Early career researchers in HDS need to develop these skills during their postdoctoral or doctorate research journey. Therefore, two participants suggested those concepts to be included in HDS courses. For example, one participant discussed the benefit of having training on high-performance computing:

> *“[…] having more courses that teach the basics of high-performance computing will be very useful (P7).”*

In summary, course materials were discussed a total of 14 times by all participants. The results indicate that visual materials were more favoured by participants (referenced 8 times by 6 participants), although they also acknowledged the benefits of textual materials (referenced 4 times by 4 participants). Also, participants highlighted the importance of including instruction on GitHub, high-performance computing, and Linux, which are essential to know during doctoral and postdoctoral research in HDS.

#### P2: Course length and level

Course length and level aspects were discussed by 3 participants 5 times. The field of HDS involves learners with diverse levels of computational skills as well as knowledge in biology. Therefore, course designers should carefully consider the level of course materials to avoid making them too easy or too difficult for students. Course duration is also important, as a course that is either too short or too lengthy may fail to engage students.

For instance, a master’s student with biology background expressed a desire for a more extensive coding course, stating:

> *“I think I do wish that we had more, because at the start, we only had a very short course of coding. I wish that was a bit longer(P10).”*

A PhD student with computational background mentioned a course design that overwhelmed them due to an excessive amount of content:

> *“[…] For example, some lecturers, they put a lot of information into the course and students find it very difficult to follow all the content (P6).”*

A PhD student highlighted the impact of teaching speed on the learning experience:

> *“We had the professor typing up the code, and we all had our individual laptops, and we were all following in theory. That sounds like it’s a really good idea because everyone’s doing it together, but they went way too fast for us, and everyone ends up being in different parts of the material. I think it’s a good idea still. I think you kind of have to think about how much you’re going to fit in one session. Right? Because if you’re going through it so quickly, then it’s not going to work. Then everyone’s going to be like lagging behind (P9).”*

The participant thought that although coding during class altogether was an effective teaching approach, it was too fast for them and affected their learning experience. This illustrates the importance of ensuring that the course level and length align as closely as possible with students’ levels and goals.

In summary, course length and level were mentioned by 3 participants 5 times; therefore, finding a balance in HDS courses regarding the difficulty level and duration is important although it can be challenging to get right. It is recommended that course designers and teachers gain a thorough understanding of their students’ levels in advance to tailor the course accordingly [45].

#### P3: Discussion forums

This category includes quotes about discussion forums, with five codes: **immediate support** (referenced 7 times by 6 participants), **engaging all students** (referenced twice by 2 participants), **anonymised questions** (referenced once by 1 participant), **asking basic questions** (referenced once by 1 participant), and s**ubject-based discussions** (referenced once by 1 participant).

A code that received more attention than others was i**mmediate response and support**. Participants pointed out the long waiting times for answers as one of the weaknesses of online discussion forums. For instance, a participant stated:

> *“[…] in the different discussion forum there are people not able to answer immediately. That’s the problem. So in the research field especially you are not going to wait for anybody. […] (P7).”*

Two PhD students also discussed the importance of **engaging all students**:

> *“I mean, usually if you have an issue, you start with discussion, but sometimes people do not engage. Or maybe because I’m not particular interested in that topic. So that’s the reason I feel the discussion. So, the success of online course discussions depends on the level of engagement (P3).”*

In line with the quote from *P3*, a discussion subject is important, and one other PhD student suggested **subject-based discussions** that align with the students’ interests:

> *“[…] For example, on platforms like Stack Overflow or other similar websites, discussions usually are around solving problems. If it’s a common problem, discussions tend to be more active. Everyone shares their perspectives on how to solve the problem, which creates engaging discussions. However, discussing just the course material without a specific problem to solve can be challenging for students to collaborate on (P5).”*

A postdoctoral researcher emphasised the importance of encouraging students to ask even their **basic questions**:

> *“The thing is, some people might think if they ask their question, that might seem silly or very simple. But having that said that every single question is really important, even if it might seem simple, you should ask it (P1).”*

A participant mentioned the option of asking **anonymised questions** to help engage students:

> *“[…] Maybe if students are shy, they can have an anonymized feature where people don’t reveal who they are and they can ask a question, so they don’t feel like they’re asking anything stupid or something (P7).”*

In summary, weaknesses of online discussion forums and suggestions for improvement were discussed by early career researchers (12 times by 7 participants). It is essential to improve discussion forums in online health data science courses to provide students with immediate support and responses, as it was pointed out frequently by participants (7 times by 6 participants). Allowing for anonymised questions can encourage students to ask even basic questions. Teachers could encourage all learners to participate in discussions by initiating subject-based discussions that motivate students, even to ask simple questions. Additionally, early career researchers found value in meeting and discussing concepts with their colleagues to make sure they understood concepts correctly.

#### P4: Teaching method

The teaching approach, referenced 43 times by all participants, includes six codes: **teaching through discussion** (referenced 15 times by 8 participants), **lecture before discussion** (referenced 1 time by one participant), **practical emphasis** (referenced 17 times by 8 participants), **theoretical emphasis** (referenced 2 times by 2 participants), **teaching with simple language** (referenced 3 times by one participant), and **having interdisciplinary lecture** (referenced 2 times by one participant).

**Teaching through discussion** was referenced 15 times by 8 participants who expressed a preference for interactive teaching over lecture-based teaching. For example, one master’s student highlighted this preference by stating:

> *“I like it when we’re all trying to work on one problem and then talking about it and solving it together rather than lecturing teaching (P10).”*

Conversely, another PhD student preferred having **lectures before discussions**:

> *“Before going for the discussion, it is very important to understand concepts. If you are not understood concepts and going for the discussion it is very difficult […] (P4).”*

Participants also shared their preferences for teaching with a focus on **practical** concepts over theoretical ones. The majority (8 of all participants) favoured an approach that emphasises practical aspects and involves hands-on experiences. For instance, the postdoctoral researcher stated:

> *“If it is more coding or computing way of the courses, then I would suggest having hands-on experience with coding. If it is more on the biological side, I also suggest having some hands-on experience of the biological (P1).”*

However, two students argued that theoretical aspects are foundational and should precede practical applications:

> *“I think you need to have some theory and then apply to a practical part because otherwise it’s like just theory and you don’t know how to be a bit more flexible to apply and then actually come with the problem that arises when you face the situation (P8).”*

A postdoctoral researcher also expressed a preference for t**eaching with simple language**:

> *“The video lectures that have very simple cartoon things in it that makes everything more simple so that all the audiences with every level can understand that […] I should say then simplified videos (P1).”*

This participant also discussed the benefits of **interdisciplinary lectures**, suggesting that lecturers from various backgrounds, such as biology and computer science, could prepare students for interdisciplinary environments:

> *“For me if during my PhD or the first few days of the postdoc, if there was a biologist who came to our sessions and talk about that computational problem from a biologist perspective, it’s like an inviting lecturer that for example, you have a lecturer that is learning you how to do the statistical analysis on the biological data. And you have, for example, twelve sessions that a computational person is teaching you. Second, you might have an introduction of how clinical people are talking, what their lang uage is, and this can really help if it is happening in the earlier stages in their academic life (P1).”*

In summary, discussion-based teaching (referenced 15 times by 8 participants) was more favourable than lecture-dominated courses in health data science education. It provides students with the opportunity to engage in meaningful discussions with their peers, enhancing their learning experience. Therefore, incorporating a substantial amount of constructive discussion-based teaching or interactive lectures may enhance health data science education. Participants also emphasised their preference for hands-on experience with a focus on practical aspects (referenced 17 times by 8 participants). Consequently, incorporating real-world projects and explaining concepts within the context of real-world problems, having lecturers come from diverse backgrounds, and explaining technical concepts using simple language, can benefit health data science education.

## 4 Discussion

The most frequently discussed learning strategy was peer learning and help seeking (**S1: Collaboration**). This learning strategy is aligned with well-recognised learning strategies in education [11, 46]. Research has shown that this strategy is effective in improving student learning experience and performance [11, 32, 46, 47]. Seeking help from those who have navigated a similar path can help learners overcome difficulties [48]. Engaging in group discussions and asking questions in interdisciplinary settings not only facilitates knowledge sharing across different fields but also enhances interdisciplinary communication skills [48]. Students with backgrounds in biology can provide insights into biological questions, while those with a computational background can assist their peers in gaining a deeper understanding of statistics and programming. Pair programming can also support collaborative learning [49], and it has been successfully employed in an HDS education setting [50]. Additionally, meeting one-on-one with supervisors can be a valuable resource for addressing questions [51].

Information seeking [32, 33] (**S2: Information seeking**) was the second most frequently discussed learning strategy. This learning strategy involves seeking information by participating in courses and conferences, or reading books and papers to address the lack of foundational knowledge [2]. According to our findings, it can be inferred that students’ goals have impact on their approach in seeking information. We found that HDS learners used goal orientation strategy (**S8**) for a course or research work and then looked at the learning materials to prioritise key concepts essential to achieve their learning goals. By using the goal orientation strategy, students can decide whether they would like to explore topics beyond syllabus, if this aligns with their goals and time constraints. Goal-setting theory [51], which is based on well-known psychological research, demonstrates that having goals can help students to perform effectively and focus more than others on their performance. Also, learners who have a goal are more likely to achieve better performance, go beyond syllabus and course standard expectation, using strategies necessary to regulate their attention and effort [11, 36, 51–53]. However, according to **(S7)** early-career researchers in HDS believed it is not always necessary to focus on every detail; focusing on acquiring the necessary discipline-specific knowledge could be enough, depending on a student’s aims.

Actively engaging (**S3: Active learning**) in an actual HDS project and acquiring skills through problem-solving can be an effective learning strategy [34]. Previous research showed that active learning strategies can help students to achieve better outcomes [52]. One of the strategies to actively engage in HDS is practice in coding, which can help improve programming skills [48]. Also, reviewing sample code and previous projects can aid in overcoming challenges in learning programming and developing computational skills [53–55].

Providing students with a variety of resources such as videos and textbooks is important, but enhancing reading materials and video lectures with visually appealing elements like diagrams and flowcharts can aid comprehension [56]. Aligning course content with the current needs of HDS students and including instructions on technical tools such as GitHub and HPC is valuable (**P1: Course materials**) [57]. Given the fact that students come from diverse backgrounds and skill levels, it is essential to maintain a balanced pace and level in a course [57–60].

Encouraging students to engage by asking questions and encouraging peer discussions during lectures can improve students’ understanding (**P4: Teaching method**) [34]. Early-career researchers expressed a preference for interactive lectures that allow them to engage in discussions rather than passively receiving information. Previous research also emphasised having interactive and project-based lectures for health data science [57, 61, 62]. Presenting concepts by working on a real project step by step is more favourable for students than explaining theories alone (**P4: Teaching method**). Students appreciate seeing how they can apply their knowledge to solve real HDS problems. Also, having teachers from both biology and computer science backgrounds to explain different course topics not only prepares students for the interdisciplinary nature of HDS but also addresses the challenge of teacher knowledge gaps (**P4: Teaching method**).

Finally, while online courses offer accessibility for remote learners, there is room for improvement in online discussion forums (**P3: Discussion forums**) [63]. Suggestions for improvement include providing options for anonymous questioning, offering prompt responses, having subject for a discussion, and encouraging all students to ask questions, even basic ones.

The results of this study, along with the suggestions derived from these findings, have the potential to raise learners’ awareness of effective learning strategies for overcoming challenges. Furthermore, valuable insights from this study, such the importance of interactive teaching and on hands-on learning, can inform HDS education improvement.

## 5 Limitations and Future Work

To enhance the diversity of the data, it would be beneficial to conduct interviews with learners from around the world, as our study currently only focuses on participants from UK institutions. Additionally, increasing the number of interviews might strengthen results further. Furthermore, it is worth noting that the results are only based on self-declared information, which might be consciously or unconsciously affected by different factors, such as social desirability. To strengthen findings, it would be advantageous to complement this qualitative analysis with quantitative approaches. Analysing click - stream data of student behaviour in HDS courses could provide a more rigorous basis for research findings. In our study, we focused exclusively on suggestions from the learners’ perspective. Conducting interviews with teachers and lecturers could offer a complementary perspective and uncover additional insights into HDS education. In future research, we plan to incorporate quantitative analysis and compare the findings from qualitative and quantitative approaches to provide more robust insights and recommendations.

## 6 Conclusions

We conducted a qualitative content analysis to shed light on learning strategies of early-career researchers in health data science. We also provided insight into effective teaching approaches and course designs for HDS. Our results, based on interviews with early career researchers in two UK institutions, reveal that learners employ various strategies, such as peer learning and help seeking, information seeking, and actively practising coding and problem solving. Participants expressed a preference for interactive and practical lectures that provide opportunities to discuss real-world problems and collaboratively solve them within the classroom. Additionally, improvements in online course discussions are needed, with an emphasis on providing prompt responses to encourage engagement.

## List of Abbreviations

AI: Artificial Intelligence
HI: Health Data Science
MSLQ: Motivated Strategies for Learning Questionnaire
P: Participant
RQ: Research Question
SRQR: Standards for Reporting Qualitative Research

## Declarations

## Ethical approval and consent to participate

This research has been approved by the ethics committee of the School of Informatics, University of Edinburgh [Reference number: #88883]. Necessary ethical considerations were performed before, during, and after conducting the research. The participation in the study was voluntary, and necessary information about the study was sent to participants before the interview. Informed consent was obtained from all participants. Participants signed a consent form before starting the interview.

## Consent for publication

Not applicable.

## Availability of data and materials

The whole dataset used during the current study is not publicly available due to ethical restrictions. However, the processed data and supporting data is reported in the manuscript. To get access to the whole data, an ethical approval from the Ethical Committee of the University of Edinburgh is needed.

## Authors’ contributions

All authors designed and conceptualised the study. NR conducted the interviews under the supervision of AM, and MG. NR and MG iteratively carried out the coding process. NR wrote the manuscript, which was enhanced by AM, MG, and KG. The final version of the manuscript was reviewed and accepted by all authors.

## Supplementary materials

**Additional file 1:** SRQR form

**Additional file 2:** Interview questions

**Additional file 3:** Consent and participation information forms

## 7 Acknowledgement

We would like to thank all the participants who generously shared valuable insights about their learning experiences in HDS. Also, we extend our thanks to Dr. Syed Haider and Prof. Claudio Angione for their invaluable assistance in recruiting participants. We also thank Manataki lab members for their useful comments that helped in improving the readability of the manuscript.

## References

1. Kolachalama, V.B. and P.S. Garg, Machine learning and medical education. NPJ digital medicine, 2018. 1(1): p. 54.

2. Işık, E.B., et al., Grand challenges in bioinformatics education and training. Nature Biotechnology, 2023. 41(8): p. 1171–1174.

3. Wan, T. and V. Gurupur, Understanding the difference between healthcare informatics and healthcare data analytics in the present state of health care management. Health services research and managerial epidemiology, 2020. 7: p. 2333392820952668.

4. Rohani, N., et al. Early prediction of student performance in a health data science MOOC. in Proceedings of the 16th International Conference on Educational Data Mining. 2023. International Educational Data Mining Society.

5. Executive summary of the meeting of the 2020 ashp Commission on goals: preparing the healthcare workforce for a digital future. American Journal of Health-System Pharmacy, 2020. 78(5): p. 447–453.

6. Seth, P., et al., Data science as a core competency in undergraduate medical education in the age of artificial intelligence in health care. JMIR medical education, 2023. 9(1): p. e46344.

7. Grunhut, J., A.T. Wyatt, and O. Marques, Educating Future Physicians in Artificial Intelligence (AI): An Integrative Review and Proposed Changes. Journal of Medical Education and Curricular Development, 2021. 8: p. 23821205211036836.

8. Magana, A.J., et al., A survey of scholarly literature describing the field of bioinformatics education and bioinformatics educational research. CBE— Life Sciences Education, 2014. 13(4): p. 607–623.

9. Matcha, W., et al., Analytics of Learning Strategies: Role of Course Design and Delivery Modality. Journal of Learning Analytics, 2020. 7(2): p. 45–71.

10. Mazzetti, G., et al., The impact of learning strategies and future orientation on academic success: The moderating role of academic self-efficacy among Italian undergraduate students. Education Sciences, 2020. 10(5): p. 134.

11. Pintrich, P.R., A manual for the use of the Motivated Strategies for Learning Questionnaire (MSLQ). 1991.

12. Weinstein, C.E. and V.L. Underwood, Learning strategies: The how of learning, in Thinking and learning skills. 2014, Routledge. p. 241–258.

13. Bickerdike, A., et al., Learning strategies, study habits and social networking activity of undergraduate medical students. International journal of medical education, 2016. 7: p. 230.

14. Stark, J.S., et al., Disciplinary differences in course planning. The Review of Higher Education, 1990. 13(2): p. 141–165.

15. Schuster, D., et al., Learning of core disciplinary ideas: Efficacy comparison of two contrasting modes of science instruction. Research in Science Education, 2018. 48: p. 389–435.

16. et al., How approaches to teaching are affected by discipline and teaching context. Studies in Higher education, 2006. 31(03): p. 285–298.

17. Cameron, L., How learning designs, teaching methods and activities differ by discipline in Australian universities. Journal of learning design, 2017. 10: p. 69–84.

18. Stark, J.S., Planning introductory college courses: Content, context and form. Instructional Science, 2000. 28: p. 413–438.

19. Entwistle, N. and H. Tait, Approaches to studying and perceptions of the learning environment across disciplines. New directions for teaching and learning, 1995. 1995(64): p. 93–103.

20. Danielson, C., Enhancing professional practice: A framework for teaching. 2007: AsCD.

21. O’Brien, B.C., et al., Standards for reporting qualitative research: a synthesis of recommendations. Academic medicine, 2014. 89(9): p. 1245–1251.

22. Rohani, N., et al., Providing Insights into Health Data Science Education through Artificial Intelligence. bioRxiv, 2024: p. 2024.03.22.586308.

23. Ng, S.L., et al., Qualitative research in medical education: methodologies and methods. Understanding medical education: Evidence, theory, and practice, 2018: p. 427–441.

24. Hsieh, H.-F. and S.E. Shannon, Three approaches to qualitative content analysis. Qualitative health research, 2005. 15(9): p. 1277–1288.

25. Elo, S. and H. Kyngäs, The qualitative content analysis process. Journal of advanced nursing, 2008. 62(1): p. 107–115.

26. Wolters, C.A., L.Y. Shirley, and P.R. Pintrich, The relation between goal orientation and students’ motivational beliefs and self-regulated learning. Learning and individual differences, 1996. 8(3): p. 211–238.

27. Winne, P., Self-regulated learning. SFU Educational Review, 2016. 9.

28. Biggs, J.B., Student Approaches to Learning and Studying. Research Monograph. 1987: ERIC.

29. AssemblyAI. AssemblyAI’s Python SDK. 2023; Available from: https://www.assemblyai.com/.

30. Dhakal, K., NVivo. Journal of the Medical Library Association: JMLA, 2022. 110(2): p. 270.

31. Abrahamsson, S. and M. Dávila López, Comparison of online learning designs during the COVID-19 pandemic within bioinformatics courses in higher education. Bioinformatics, 2021. 37(Supplement_1): p. i9–i15.

32. Schunk, D.H., Social cognitive theory and self-regulated learning, in Self-regulated learning and academic achievement. 2013, Routledge. p. 119–144.

33. Nandagopal, K. and K.A. Ericsson, An expert performance approach to the study of individual differences in self-regulated learning activities in upper-level college students. Learning and Individual Differences, 2012. 22(5): p. 597–609.

34. Prince, M., Does active learning work? A review of the research. Journal of engineering education, 2004. 93(3): p. 223–231.

35. Albanese, M.A. and L.C. Dast, Problem-based learning. Understanding medical education: Evidence, theory and practice, 2013: p. 61–79.

36. Dostál, J., Theory of problem solving. Procedia-Social and Behavioral Sciences, 2015. 174: p. 2798–2805.

37. Duncan, T.G. and W.J. McKeachie, The making of the motivated strategies for learning questionnaire. Educational psychologist, 2005. 40(2): p. 117–128.

38. Soloman, B.A. and R.M. Felder, Index of learning styles questionnaire. NC State University. Available online at: http://www.engr.ncsu.edu/learningstyles/ilsweb.html (last visited on 14.05. 2010), 2005. 70.

39. Felder, R.M. and J. Spurlin, Index of learning styles. International Journal of Engineering Education, 1991.

40. Gregorc, A.F. and K.A. Butler, Learning is a matter of style. VocEd, 1984. 59(3): p. 27–29.

41. Floyd, K.S., S. Harrington, and J. Santiago, The effect of engagement and perceived course value on deep and surface learning strategies. Informing Science, 2009. 12: p. 181.

42. Postareff, L., A. Parpala, and S. Lindblom-Ylänne, Factors contributing to changes in a deep approach to learning in different learning environments. Learning Environments Research, 2015. 18: p. 315–333.

43. Marton, F. and R. Säljö, On qualitative differences in learning: I—Outcome and process. British journal of educational psychology, 1976. 46(1): p. 4–11.

44. Karagiannopoulou, E. and N. Entwistle, Students’ learning characteristics, perceptions of small-Group University teaching, and understanding through a “meeting of minds”. Frontiers in psychology, 2019. 10: p. 444.

45. Gross, L.J., Points of view: the interface of mathematics and biology: interdisciplinarity and the undergraduate biology curriculum: finding a balance. Cell Biology Education, 2004. 3(2): p. 85–87.

46. Newman, R.S., Adaptive help seeking: A strategy of self-regulated learning, in Self-regulation of learning and performance. 2023, Routledge. p. 283–301.

47. Coliñir, J.H., et al., Characteristics and impacts of peer assisted learning in university studies in health science: A systematic review. Revista Clínica Española (English Edition), 2022. 222(1): p. 44–53.

48. Carey, M.A. and J.A. Papin, Ten simple rules for biologists learning to program. 2018, Public Library of Science. p. e1005871.

49. Salleh, N., E. Mendes, and J. Grundy, Empirical Studies of Pair Programming for CS/SE Teaching in Higher Education: A Systematic Literature Review. IEEE Transactions on Software Engineering, 2011. 37(4): p. 509–525.

50. Doudesis, D. and A. Manataki, Data science in undergraduate medicine: Course overview and student perspectives. International Journal of Medical Informatics, 2022. 159: p. 104668.

51. Almusaed, A. and A. Almssad, The Role of the Supervisor on Developing PhD Students’ Skills. International Society for Technology, Education, and Science, 2020.

52. Aji, C.A. and M.J. Khan, The impact of active learning on students’ academic performance. Open Journal of Social Sciences, 2019. 7(03).

53. Van Merrienboer, J.J. and H.P. Krammer, Instructional strategies and tactics for the design of introductory computer programming courses in high school. Instructional Science, 1987. 16: p. 251–285.

54. Busjahn, T. and C. Schulte. The use of code reading in teaching programming. in Proceedings of the 13th Koli Calling international conference on computing education research. 2013.

55. Van Gog, T. and N. Rummel, Example-based learning: Integrating cognitive and social-cognitive research perspectives. Educational psychology review, 2010. 22: p. 155–174.

56. Chen, C.-M. and Y.-C. Sun, Assessing the effects of different multimedia materials on emotions and learning performance for visual and verbal style learners. Computers & Education, 2012. 59(4): p. 1273–1285.

57. Via, A., et al., Best practices in bioinformatics training for life scientists. Briefings in bioinformatics, 2013. 14(5): p. 528–537.

58. Ranganathan, S., Bioinformatics education—perspectives and challenges. PLoS computational biology, 2005. 1(6): p. e52.

59. Tastan Bishop, Ö., et al., Bioinformatics education—perspectives and challenges out of Africa. Briefings in bioinformatics, 2015. 16(2): p. 355–364.

60. Schneider, M.V., et al., Bioinformatics training: a review of challenges, actions and support requirements. Briefings in bioinformatics, 2010. 11(6): p. 544–551.

61. Emery, L.R. and S.L. Morgan, The application of project-based learning in bioinformatics training. PLoS computational biology, 2017. 13(8): p. e1005620.

62. Eglen, S., A Quick Guide to Teaching R Programming to Computational Biology Students. Retrieved February 20, 2016. 2009.

63. de Lima, D.P., et al., What to expect, and how to improve online discussion forums: the instructors’ perspective. Journal of Internet Services and Applications, 2019. 10: p. 1–15.

