## Additional file1 for "Insights into Health Data Science Education: A Qualitative Content Analysis"

### Standards for Reporting Qualitative Research (SRQR)*

http://www.equator-network.org/reporting-guidelines/srqr/

**Title and abstract** **Page/line no(s).**

| | |
|---|---|
| **Title** - Concise description of the nature and topic of the study Identifying the study as qualitative or indicating the approach (e.g., ethnography, grounded theory) or data collection methods (e.g., interview, focus group) is recommended | Page 1 |
| **Abstract** - Summary of key elements of the study using the abstract format of the intended publication; typically includes background, purpose, methods, results, and conclusions | Page1 |

**Introduction**

| | |
|---|---|
| **Problem formulation** - Description and significance of the problem/phenomenon studied; review of relevant theory and empirical work; problem statement | Page 2-3 |
| **Purpose or research question** - Purpose of the study and specific objectives or questions | Page 3 |

**Methods**

| | |
|---|---|
| **Qualitative approach and research paradigm** - Qualitative approach (e.g., ethnography, grounded theory, case study, phenomenology, narrative research) and guiding theory if appropriate; identifying the research paradigm (e.g., postpositivist, constructivist/ interpretivist) is also recommended; rationale** | Page 4 |
| **Researcher characteristics and reflexivity** - Researchers' characteristics that may influence the research, including personal attributes, qualifications/experience, relationship with participants, assumptions, and/or presuppositions; potential or actual interaction between researchers' characteristics and the research questions, approach, methods, results, and/or transferability | Page 6 |
| **Context** - Setting/site and salient contextual factors; rationale** | Page 4-5 |
| **Sampling strategy** - How and why research participants, documents, or events were selected; criteria for deciding when no further sampling was necessary (e.g., sampling saturation); rationale** | Page 4-6 |
| **Ethical issues pertaining to human subjects** - Documentation of approval by an appropriate ethics review board and participant consent, or explanation for lack thereof; other confidentiality and data security issues | Page 5, 23 |
| **Data collection methods** - Types of data collected; details of data collection procedures including (as appropriate) start and stop dates of data collection and analysis, iterative process, triangulation of sources/methods, and modification of procedures in response to evolving study findings; rationale** | Page 4-5 |

| | |
|---|---|
| **Data collection instruments and technologies** - Description of instruments (e.g., interview guides, questionnaires) and devices (e.g., audio recorders) used for data collection; if/how the instrument(s) changed over the course of the study | Page 6 |
| **Units of study** - Number and relevant characteristics of participants, documents, or events included in the study; level of participation (could be reported in results) | Page 5 |
| **Data processing** - Methods for processing data prior to and during analysis, including transcription, data entry, data management and security, verification of data integrity, data coding, and anonymization/de-identification of excerpts | Page 4-6 |
| **Data analysis** - Process by which inferences, themes, etc., were identified and developed, including the researchers involved in data analysis; usually references a specific paradigm or approach; rationale** | Page 4-6 |
| **Techniques to enhance trustworthiness** - Techniques to enhance trustworthiness and credibility of data analysis (e.g., member checking, audit trail, triangulation); rationale** | Page 4-6 |

**Results/findings**

| | |
|---|---|
| **Synthesis and interpretation** - Main findings (e.g., interpretations, inferences, and themes); might include development of a theory or model, or integration with prior research or theory | Page 6-21 |
| **Links to empirical data** - Evidence (e.g., quotes, field notes, text excerpts, photographs) to substantiate analytic findings | Page 6-21, Table 1 and Table 2 |

**Discussion**

| | |
|---|---|
| **Integration with prior work, implications, transferability, and contribution(s) to the field -** Short summary of main findings; explanation of how findings and conclusions connect to, support, elaborate on, or challenge conclusions of earlier scholarship; discussion of scope of application/generalizability; identification of unique contribution(s) to scholarship in a discipline or field | Page 21-22 |
| **Limitations** - Trustworthiness and limitations of findings | Page 22 |

**Other**

| | |
|---|---|
| **Conflicts of interest** - Potential sources of influence or perceived influence on study conduct and conclusions; how these were managed | Page 6 |
| **Funding** - Sources of funding and other support; role of funders in data collection, interpretation, and reporting | Page 23 |

*The authors created the SRQR by searching the literature to identify guidelines, reporting standards, and critical appraisal criteria for qualitative research; reviewing the reference lists of retrieved sources; and contacting experts to gain feedback. The SRQR aims to improve the transparency of all aspects of qualitative research by providing clear standards for reporting qualitative research.

**The rationale should briefly discuss the justification for choosing that theory, approach, method, or technique rather than other options available, the assumptions and limitations implicit in those choices, and how those choices influence study conclusions and transferability. As appropriate, the rationale for several items might be discussed together.

**Reference:**

O'Brien BC, Harris IB, Beckman TJ, Reed DA, Cook DA. **Standards for reporting qualitative research: a synthesis of recommendations.** *Academic Medicine*, Vol. 89, No. 9 / Sept 2014
DOI: 10.1097/ACM.0000000000000388
