## Additional file 2 for "Insights into Health Data Science Education: A Qualitative Content Analysis"

**Interview questions**

**Aims:**

- How early-career researchers in HDS manage their educational challenges?
- What course design and teaching approach could help them to better learn HDS topics.

**Place:** private room, face to face individual interview

Time: Evening or afternoon. Based on their availability.

**Expected duration:** 20-30 minutes.

**Equipment:** Paper, pen, recording their voice, consent form, participation information will be emailed to them before the interview. They will have time to ask questions.

**Questions:**

Q1 (triggering question). Could you please provide information about your research project, in two or three sentences?

Q1.1. Is your background, for example undergraduate study, in the field of computational sciences or biological sciences?

Q2. What challenges did you encounter while you were being trained in computational methods/biological concepts?

Q3. How do you manage your learning HDS? When you enrolled in a HDS course, what approach did you adopt to optimise your learning?

Q4. What would be in your opinion the ideal design for HDS course?

Q5. What are your thoughts on incorporating discussion forums into an online course?

Q6. Do you have any ideas that could enhance the design of a HDS course?
