## Additional file 3 for "Insights into Health Data Science Education: A Qualitative Content Analysis"

and Areti Manataki [4]

[1*] Usher Institute, University of Edinburgh, UK
[2] Moray House School of Education and Sport, University of Edinburgh, UK
[3] School of Informatics, University of Edinburgh, UK
[4] School of Computer Science, University of St Andrews, UK

### Data-driven Insights into Health Data Science Training

#### Participant Information Sheet

**Introduction**

We appreciate your interest in being part of this research. This document presents an overview of the study's objective and provides a description of your involvement and rights as a participant.

**1. What is the research about?**

This research project aims to enhance students' experience in precision medicine and health data science courses. The objective of the interview is to gain insights into students' learning experiences, challenges, and strategies in courses related to health data science/precision medicine. By understanding these aspects, we can facilitate improvements in current and future course designs in the field of health data science.

**2. Why have I been invited to take part?**

You are invited to participate in this study due to your involvement within the fields associated with health data science/precision medicine.

**3. Do I have to take part?**

The decision to participate in this study is entirely yours to make. You have the freedom to change your mind and discontinue or terminate your involvement at any time, without the need to provide a justification. Opting out or choosing not to participate will not have any impact on you or the services you currently access. If you choose to withdraw from the study, we will not retain any information you have previously provided, unless you grant us permission to do so. However, it is important to note that once the study concludes (in September 2023), your data will be anonymized, making it impossible to retract or withdraw your data.

**4. What will my involvement be?**

If you choose to participate, you will receive a copy of this information sheet for your records. Subsequently, you will be requested to provide consent by signing a consent form or giving oral consent, based on your preference. You will be provided with a copy of the consent form to retain, even if you opt for oral consent. It is important to note that your oral consent will be recorded for reference. You have the freedom to express any topics you do not wish to discuss or any questions you prefer not to answer. Moreover, you can request to pause or finish the interview at any time. With your consent, certain conversations may be recorded for the purpose of ensuring accuracy.

**5. What will my information be used for?**

The information gathered will be utilized for the purpose of a PhD project aiming to enhance the comprehension of challenges in precision medicine training, students' learning strategies in precision medicine courses, and the development of an effective course design in this field. The data will be analysed as part of a PhD thesis and may potentially be incorporated into academic publications.

**6. Will my taking part and my data be kept confidential? Will it be anonymised?**

All the data gathered throughout the research process will be treated with strict confidentiality. Your information will be fully anonymized to ensure your privacy and confidentiality.

**7. How will we use information about you?**

To participate in this research project, you will be requested to provide your name and contact information, which will be accessed solely by the researcher. These details will also be utilized to share research findings with you. Rest assured, we will uphold the highest standards of data protection and security in compliance with GDPR guidelines. Any audio recordings made will be deleted once they have been transcribed. Your data will be accessible exclusively by the researcher and will be stored securely in an electronically password-protected computer file, with no paper records involved.

**8. Where can you find out more about how your information is used?**

To obtain further details on how your information will be utilized, you can inquire with Ms. Narjes Rohani at or contact the University of Edinburgh Data Protection Officer at. Please note that your anonymized data will be retained for a minimum of 10 years. After the study ended, no personally identifiable information will be retained about you. However, your anonymized data may be employed in future research projects that have received ethical approval.

**9. Will the research be audio or visually recorded?**

Interviews will be conducted with audio recording. The decision to permit this recording is entirely yours, and you will be requested to provide your consent beforehand. In the event of your agreement, all recordings will be treated as confidential and solely accessible by the researcher.

**10. What are the possible benefits of taking part?**

Participating in this research does not entail direct individual benefits. However, your involvement will significantly contribute to the overarching goal of enhancing students' learning experience in precision medicine and related fields.

**12. Who has reviewed this study?**

This research proposal has undergone rigorous scrutiny and has received ethical approval from the Ethics Committee of the School of Informatics, University of Edinburgh.

The study is being conducted under the supervision of Ms. Narjes Rohani, a PhD student at the University of Edinburgh.

Funding for this research has been provided by the Medical Research Council, with the grant number MR/N013166/1.

**15. What if I have a question?**

*For general information about how the University of Edinburgh looks after research data go to: https://www.ed.ac.uk/records-management/privacy-notice-research*

If you have any further questions regarding this study, please contact us:

**Researcher** Narjes Rohani


**Supervisors:**
Dr. Areti Manataki,

Dr. Kobi Gal

Dr. Michael Gallagher

If you are happy to take part in this study, please sign the consent sheet attached or provide oral consent.

### Participant Consent Form

| | |
|---|---|
| Project title: | Data-driven insights into precision medicine training |
| Principal investigator (PI): | Dr. Kobi Gal, Dr. Areti Manataki, Dr. Michael Gallagher |
| Researcher: | Ms. Narjes Rohani |
| PI contact details: | +44 (0) 131 651 5657 |

By participating in the study you agree that:

- I have read and understood the Participant Information Sheet for the above study, that I have had the opportunity to ask questions, and that any questions I had were answered to my satisfaction.
- My participation is voluntary, and that I can withdraw at any time without giving a reason. Withdrawing will not affect any of my rights.
- I consent to my anonymised data being used in PhD thesis, academic publications and presentations.
- I understand that my anonymised data will be stored for the duration outlined in the Participant Information Sheet.

**Please tick yes or no for each of these statements.**

**1.** I agree to being audio recorded.

| | |
|---|---|
| **Yes** | **No** |

**2.** I allow my data to be used in future studies that are ethically approved.

| | |
|---|---|
| **Yes** | **No** |

**3.** I agree to take part in this study.

| | Yes | No |
|---|---|---|

| Name of person giving consent | Date<br>dd/mm/yy | Signature |
|---|---|---|
| Name of person taking consent | Date<br>dd/mm/yy | Signature |
